# Individual difference in serial dependence results from opposite influences of perceptual choices and motor responses

**DOI:** 10.1101/631309

**Authors:** Huihui Zhang, David Alais

## Abstract

Natural image statistics exhibit temporal regularities of slow changes and short-term correlations and visual perception, too, is biased towards recently seen stimuli, i.e., a positive serial dependence. Some studies report strong individual differences in serial dependence in perceptual decision making: some observers show positive serial effects, others repulsive effects, and some show no bias. To understand these contrasting results, this study separates the influences of physical stimuli *per se*, perceptual choices and motor responses on serial dependence in perceptual decision making. In two experiments, human observers reported which orientation (45° or −45°, at threshold contrast) they perceived. Experiment 1, used a consistent mapping between stimulus and response buttons while in Experiment 2, observers did two tasks: one with a consistent stimulus-response mapping, the other with a random stimulus-response mapping (perceptual choice and motor response unrelated). Results show that the stimulus percept (not the physical stimulus *per se)* affected subsequent perceptual choices in an attractive way, and that motor responses produced a repulsive serial effect. When the choice-response mapping was consistent (inseparable choice and response, typical of most experiments), individual differences in the overall serial effect was observed: some were positive, some repulsive, and some were bias-free. These individual differences likely reflect different relative weightings in individuals of a positive choice bias and a repulsive motor bias.

## Introduction

Numerous studies have shown that current perceptual decisions can be affected by recent history. Using continuous measures of perception (reproducing the presented stimuli or rating on a scale), it has been shown that visual perception at a given moment is biased towards the recently seen stimuli (i.e., a positive serial dependence). This has been established for various attributes, from basic features including orientation (Cicchini, Mikellidou, & Burr, 2017; Fischer & Whitney, 2014; Fritsche, Mostert, & de Lange, 2017) and numerosity (Cicchini, Anobile, & Burr, 2014) to face identity (Liberman, Fischer, & Whitney, 2014). Using categorization tasks, positive serial dependence was found for attributes such as orientation (Cicchini et al. 2017; Norton, Fleming, & Daw, 2017), motion (Alais, Leung, & Van der Burg, 2017) and gender of face (Taubert, Alais, & Burr, 2016), while a repulsive serial dependence was found for motion-induced orientation (Alais et al., 2017), facial expression (Taubert et al., 2016). These studies investigated serial dependence by examining how the previous stimulus influences current perception, but another way to look at serial dependence is to examine how previous responses affect current perceptual decision making. It has been known since the early 20th century that human observers’ decisions depend on their previous choices (Fernberger, 1920). When using categorization tasks, studies of how present perceptual choices are influenced by preceding choices show a puzzling variety of results: some report positive serial effects and others report negative serial effects (Abrahamyan, Silva, Dakin, Carandini, & Gardner, 2016; Braun, Urai, & Donner, 2018; Fründ, Wichmann, & Macke, 2014; Raviv, Ahissar, & Loewenstein, 2012). Understanding this contradiction is the aim of the current study.

Although serial dependence has been studied extensively, what specifically is carried over from one trial to the next still remains a key question. In a typical perceptual experiment, stimulus (sensation), percept, choice and even motor response are highly correlated and thus hard to separate. Any of these processing stages could be the source of serial dependence, or there could be a cascade of serial dependencies at each stage, with the output reflecting the combination of all of them. It has been shown that a positive serial dependence is observed for briefly presented stimuli but a negative (repulsive) serial dependence is observed when the previous stimuli are presented for longer duration (adaptation-like effect). This suggests a potential repulsive contribution from sensation into overall serial dependence (Fischer & Whitney, 2014). Some studies have tried to determine whether the serial effect is due to stimulus *per se* or response *per se*. Fründ et al. (2014) modeled human observers’ performance in detecting luminance increments of differing intensities, showing that the previous response instead of previous stimulus affected current decision and that the current response was repelled away from the previous response (a repulsive bias). Using an orientation discrimination (45° or −45°) task, St John-Saaltink et al. (2016) found the same positive serial dependence on both correct choices and wrong choices, suggesting that it is the percept rather than stimulus *per se* that affects perceptual choices. Both studies show that the response is carried over trial-by-trial although the direction of bias was different in each case.

In classic binary forced-choice tasks a consistent stimulus-response mapping is used and individual differences regarding the serial dependence on previous choice/response are observed (Abrahamyan et al., 2016; Braun et al., 2018; Fründ et al. 2014). In one study, when observers were given feedback about ‘right’ or ‘wrong’ responses, some exhibited a ‘success-stay’ bias while others showed a ‘fail-switch’ bias (Abrahamyan et al., 2016). The success-stay/fail-switch strategy is advantageous in cooperative behavior (Nowak and Sigmund, 1993) and human observers seem to have different sensitivities to success and failure when applying this strategy to the task. However, individual differences are still observed even when no feedback provided (Braun et al., 2018), with some observers exhibiting a positive bias (repetition) and others showing a repulsive bias (alternation). Thus, observers seem to have different inherent (not only strategic) biases. Are individual differences a potential cause of the variously positive and negative biases found in a number of different tasks (Fründ et al., 2014; Taubert et al., 2016)? The relatively small sample of participants in those studies may result in significant positive or negative biases depending on what kind of participants are recruited.

In most studies, perceptual choices are inseparable from motor responses because the choice-response contingency remains consistent. Choosing between alternative choices can also be regarded as choosing between alternative actions. The decision-making process includes interpreting sensory information, making choices, and executing a response. Action used to be viewed as the output stage after the decision was finalized (Gold & Shadlen, 2007). However, recent evidence showing that activities in motor cortex reflect competing responses before choice commitment has challenged this view of sequential processing (Cisek & Kalaska, 2005; Klaes, Westendorff, Chakrabarti, & Gail, 2011; Pastor-Bernier & Cisek, 2011). Moreover, in our previous study, we found a behavioural oscillation of ∼10 Hz for motor response bias in perceptual decision making (Zhang, Morrone, & Alais, 2019). All these findings support an action-based decision-making theory, wherein perceptual evaluation of sensory evidence and movement planning are parallel (Cisek & Kalaska, 2010; Wispinski, Gallivan, & Chapman, 2018). If motor responses do not necessarily reflect perceptual choices, it is not clear whether the serial dependence on the previous response observed in many studies is an action-independent effect or a motor-related bias. In fact, when a trial-by-trial random response cue is used to indicate the stimulus-response mapping and thus decorrelate the perceptual choice and motor response, a positive correlation was found between previous and current perceptual choices, but an alternation bias was revealed for motor response (Pape, Noury, & Siegel, 2017; Pape & Siegel, 2016).

We addressed three questions in current study. First, is orientation discrimination at threshold (75% correct) dependent on recent history? If so, is it the stimulus or response that influences current perception? Second, can we observe individual differences in serial dependence for orientation discrimination at threshold? Third, if previous response is found to affect current perception, is the effect due to perceptual choice or motor response? Do serial dependences from perceptual choice and motor response operate in opposite directions and induce individual differences in the overall serial effect? To address these questions, we took advantage of the large dataset of 55,000 trials (N = 29, 1920 trials per participant) from our recently published study (Zhang et al., 2019), described here as Experiment 1 and which used a consistent stimulus-response mapping, and conducted a new Experiment 2 which employed both consistent and random stimulus-response mappings. To preview the results, we found that the response instead of stimulus influenced subsequent perception. There were individual differences in the sign of the serial effect (positive or repulsive bias) when using the consistent stimulus-response mapping. With a random stimulus-response mapping, there was a significant positive serial bias for choice in all participants, and a significant repulsive bias for motor responses.

## Methods

Experiment 1 involves a reanalysis of the vast amount of data collected for the experiment reported in a recently published paper (Zhang et al., 2019). The reanalysis will examine serial dependence in the orientation discrimination task used in that experiment. Experiment 2 is a new one following up results from the serial dependence analysis.

### Participants

Twenty-nine students (7 male) from the University of Sydney, aged 18-35 years, participated in Experiment 1, and all of them were naive to the purpose of the experiment. Twenty-seven new students (13 male) from the University of Sydney, aged 18-32 years, participated in Experiment 2, 24 of whom were naive to the purpose of the experiment. All the participants from Experiment 1 and 2 had normal or corrected-to-normal vision and normal audition. The study was approved by the Ethics Committee of the University of Sydney, and it was carried out in accordance with the Declaration of Helsinki. Participants gave informed consent before commencing the experiments.

### Apparatus

The apparatus is the same as used in our published study of behavioural oscillations (Zhang et al., 2019). For readability, it is described here again as follows. The experiment was run in a dimly lit room (ambient luminance, 2.1 cd/m^2)^. A PROPixx color projector (VPixx Technologies Inc.) was used to present visual stimuli on a matte white PVC screen (Epson ELP-SC21B, 1771 × 996 mm) with a resolution of 1920 × 1080, a frame rate of 120 Hz and it cast an area of 117 × 66 cm (45.4° × 26.5° of visual angle). The projector was set to quadrant mode, thereby resulting in a resolution of 960 × 540 pixels and a frame rate of 480 Hz when displaying images. The luminance output of the projector was linearized. Participants’ heads were maintained as stationary by using a chin-and-forehead rest at a viewing distance of 1.4 m. The sound stimuli were delivered bilaterally through headphones (Sennheiser HD 380 pro). All the experimental programs were developed with Matlab 2015a (MathWorks Inc., Natick, MA) and Psychophysics Toolbox.

### Stimuli and experimental procedure

The essential points concerning stimuli and procedure for Experiment 1 are summarised here and full details can be found in the Methods section of Zhang et al. (2019). The new experiment is described in full.

In both experiments, participants were required to discriminate two orientations (45 clockwise or anticlockwise, i.e., 45° or −45°) of a grating by clicking mouse buttons (Figure 1). The grating spatial frequency was 2.5 cpd and it was embedded in additive white noise and then multiplied by a Gaussian annulus window which peaked 1.0° away from the central cross and had a standard deviation of 0.3°. The white noise was randomly generated on each trial, was constant in contrast (30%) and was filtered to make its spatial frequency match the grating’s spatial frequency. The target was presented on a grey background (92.7 cd/m^2^) for a duration of 6.3 ms (3 video frames). Participants pressed a green button on the RESPONSEPixx (VPixx Technologies Inc.) using their left thumbs to initiate a trial and maintained their fixation on a central cross (0.35° wide) throughout. The target was presented 0-800 ms after the button-press after which participants reported which orientation they perceived by clicking one of two mouse buttons with their right hands.

**Figure 1.**
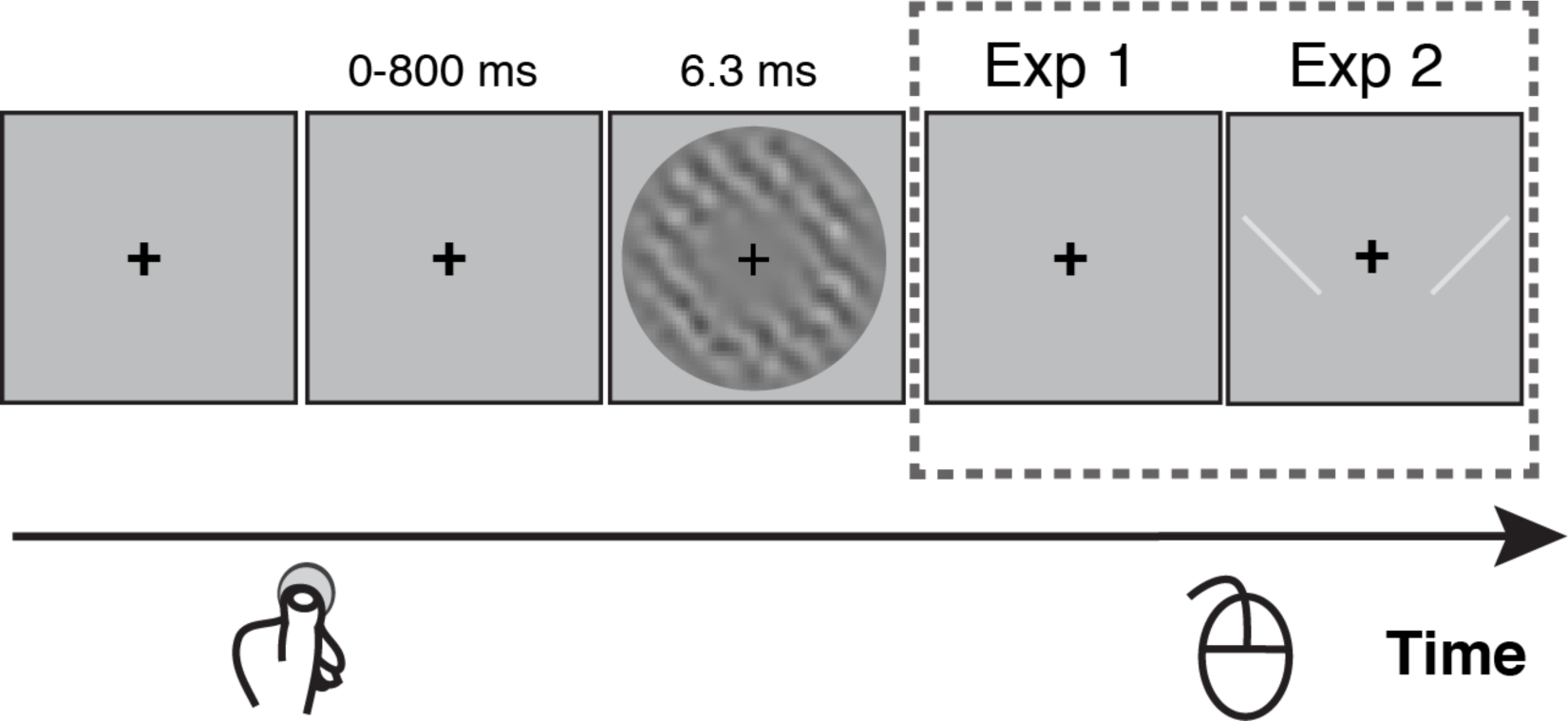
Illustration of the procedure for both experiments. Participants fixated the central cross during trials which were self-initiated by the participant by a voluntary key-press to begin each trial. After a variable time (0-800 ms), a grating (+45° or −45°) embedded in noise was presented for 6.3 ms around the fixation cross. Noise contrast was constant at 30% but grating contrast was varied to maintain threshold-level discrimination performance. Participants indicated which grating orientation (45° or −45°) they perceived using a two-button mouse. In Experiment 1, there was no response cue and the orientation-button mapping (either left click for −45° and right click for 45°, or the reverse order) was consistent for each participant but counterbalanced across participants. In Experiment 2, a visual response cue was presented after the target (two lines, 45° and −45°, either side of fixation) and remained until participants responded. In the first task, the order of the cue lines was random on each trial (either as shown, or left-right flipped). In the second task, the response cue order was the same for each participant but was counterbalanced across participants. Participants indicated orientation by choosing the location of the line that matched the grating and pressing the corresponding (left or right) mouse button.

### Experiment 1 response task

In Experiment 1, the mapping between the stimuli and mouse buttons was consistent for each participant throughout the experiment but counterbalanced across participants. Fifteen participants used an anticlockwise-left, clockwise-right mapping; while the remaining 14 used the opposite mapping (anticlockwise-right, clockwise-left).

### Experiment 2 response tasks

In Experiment 2, the mapping between stimuli and response buttons was indicated by a cue which appeared on the screen after the offset of target (see Figure 1) and remained until participants responded. The cue comprised two bright lines (±45° from vertical, 40% higher contrast than the background) presented on the left and right sides of the fixation cross (4.5° away from the central cross). The lines were multiplied by a 2D Gaussian (standard deviation = 0.3°) to soften the sharp luminance change when they were presented. Participants were required to respond to the location of the line in the cue (left or right) whose orientation matched the target grating’s orientation. For example, if the perceived orientation of the target was clockwise (45°), and the response cue showed a 45° line on the left side and a −45° line on the right side, the correct answer would be a left click. Experiment 2 manipulated the mapping between stimuli and response buttons by contrasting two response tasks. One task was as in Experiment 1: the stimulus/response mapping remained the same throughout the experiment for each participant but the left/right order (45°/−45° or −45°/45°) was counterbalanced across participants. In the other task, the left/right order in the response cue was randomized on every trial. All participants in Experiment 2 did both response tasks, with the randomised mapping always completed first, followed by the fixed mapping.

Participants were instructed that there was no time pressure to make a response and that the experiment was self-paced. There was no feedback regarding whether their response was correct or not and they were required to wait at least 2 s before pressing the button to start next trial. If they pressed the button too early, they would hear a brief beep (1000 Hz, 20 ms) and they waited two more seconds before they could initiate the trial. Before formal testing, we used an Accelerated Stochastic Approximation (ASA) procedure to adjust the contrast of the grating for each participant to yield 75% correct responses for discriminating the grating’s orientation. This contrast value was then used for the first 30 trials in the formal experiment, after which the contrast value was adjusted trial by trial using the same ASA procedure based on performance in the preceding 30 trials to ensure that performance was maintained around threshold. For Experiment 1, each participant attended two sessions over two days, each of which consisted of three blocks. Each block contained 320 trials, resulting in 1920 trials in total. Participants took a short break every 64 trials. For each task in Experiment 2, there were five blocks of 80 trials. Participants completed the two tasks in Experiment 2 consecutively in about two hours.

### Data analysis

For Experiments 1 and 2, we used signal detection theory (Green & Swets, 1966; Macmillan & Creelman; 2004) to analyse the data. In the framework of signal detection theory, both the perceptual sensitivity and criterion (decision bias) contribute to observer’s decision. Here, we are interested in how the recent past influences current perceptual decision making. If there is a bias caused by the recent history, a shift of decision criterion should be observed. Here, we chose the anticlockwise condition as ‘target’ condition and the clockwise condition as ‘noise’ condition (in a two-alternative, forced-choice context, the choice is arbitrary). That is, the hit rate is the proportion of reporting ‘anticlockwise’ orientation when the anticlockwise orientation was presented. The false alarm rate is the proportion of reporting ‘anticlockwise’ orientation when the clockwise orientation was presented. A negative value of criterion means a bias towards reporting ‘anticlockwise’ and a positive value means a bias towards ‘clockwise’. Criterion (c) was calculated using equation 1 below, where Z(HR) means the z-score of the hit rate and Z(FAR) means the z-score of the false alarm rate.

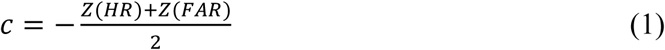

In Experiment 1, we quantified the serial dependence effect by calculating a serial dependence index (*c*_*shift*_) as shown in equation 2. The serial dependence index could be calculated based on either the previous stimulus (stimulus-based analysis) or previous choice (choice-based analysis).

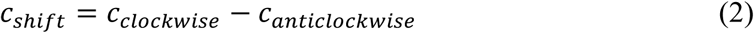

Here, the term *c*_*clockwise*_ means the criterion calculated from trials preceded by a clockwise (45°) stimulus (or a ‘clockwise’ response, for the choice-based analysis), and *c*_*anticlockwise*_ means the criterion calculated from trials preceded by an anticlockwise (−45°) stimulus (or choice). A positive *c*_*shift*_ indicates a positive serial dependence, and a negative *c*_*shift*_ indicates a repulsive serial dependence.

In Experiment 2, the inclusion of fixed and random mappings of stimuli and response buttons meant that choice and the motor response were separable. We still used equations (1) and (2) to calculate the influence of previous choices but in order to evaluate the previous response’s influence on current response, we borrowed an idea from signal detection theory. A criterion corresponding to motor bias was computed with equation (1), but the meanings of hit rate and false alarm rate were slightly different. We chose the left-click condition as the ‘target’ condition and the right-click condition as the ‘noise’ condition. That is, the hit rate is the percentage of left-clicks among trials on which the correct motor response was left-click, and the false alarm rate is the percentage of clicking left among trials on which the correct motor response was right-click. A positive criterion value means a bias towards a right click and a negative criterion means bias towards a left click. Accordingly, the serial dependence index (*c*_*shift*_) was computed with equation 3, where a positive value indicates a positive serial dependence and a negative value indicates a repulsive effect.

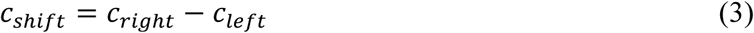

## Results

### Experiment 1

In order to examine the influence of previous choices on current perception, we need to remove the potential artefact of serial dependence caused by other factors, such as, for example, a relatively long-term preference for one choice within a block. Participants did 30 blocks in total and we calculated the bias in each block using equation 4.

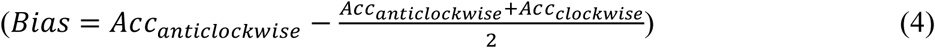

In equation 4, *Acc*_*anticlockwise*_ means the accuracy for trials presenting anticlockwise orientation, and *Acc*_*clockwise*_ means the accuracy for trials presenting clockwise orientation. The *Bias* should be zero, if there is no bias. The variance of the bias across blocks could potentially produce an artefactual serial dependence. To evaluate whether such an artefact did indeed produce a serial dependence effect, we shuffled the trial sequence in each block as this would ruin any serial dependence effect within a block but would still preserve any overall response bias within the block. Then, we calculated the serial dependence index *c*_*shift*_ with equation (2) to evaluate the influence of previous choices on current choices. We repeated this procedure 1000 times and computed the mean of the 1000 *c*_*shift*_ estimates. This mean value quantifies any artefactual serial dependence effect due to variations in response bias. We conducted this analysis for every participant, testing for artefactual serial dependence from the previous 1-5 trials.

As shown in Figure 2A, one-sample t-tests against zero revealed that there was a positive serial dependence artefact for each level of n-back (1-5) choices: *t*(28) = 3.94, 3.84, 3.73, 3.95 and 3.90, respectively, for 1-5 n-back analyses; all *ps* < 0.001. In contrast, there was no serial dependence artefact for any n-back level when serial dependence was calculated based on previous stimuli: *t*(28) = −0.024, 0.026, 0.20, 0.084 and 0.26; *ps* = 0.98, 0.98, 0.85, 0.93 and 0.80, respectively, for 1-5 n-back analyses. In addition, as shown in Figure 2B, there was a positive correlation between participants’ standard deviations across 30 blocks and their artefact of serial dependence on previous choices (*r* = 0.98, *p* < 0.001). Participants’ standard deviations across 30 blocks and their artefact of serial dependence on previous stimulus were not correlated (*r* = 0.33, *p* = 0.08). Our results show that a relatively long-term response bias can produce an artefactual serial dependence when analysing the influence of participants’ choices. Thus, for the following analyses in both Experiments 1 and 2, the artefact of serial dependence was subtracted from the original serial dependence index *c*_*shift*_ to better reveal genuine serial dependences.

**Figure 2.**
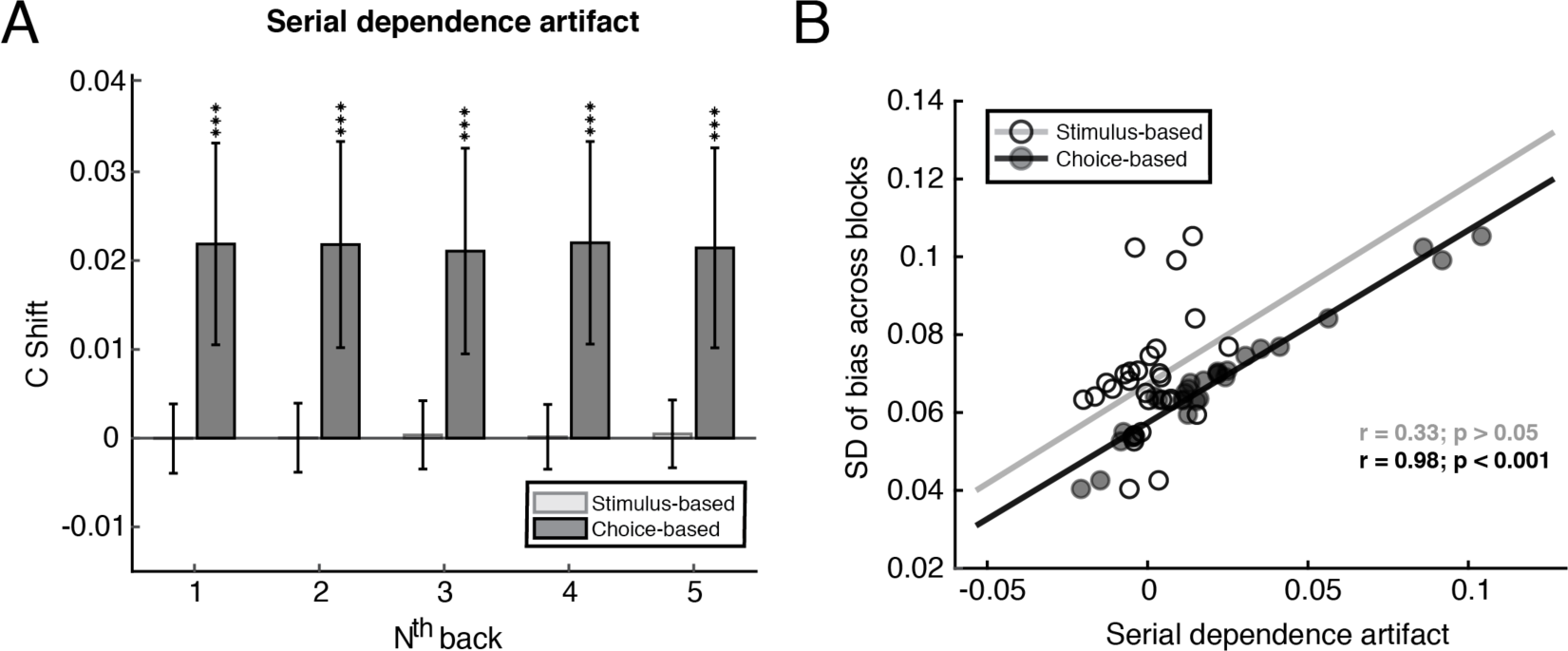
**A.** The artefact of serial dependence. The y-axis shows the serial dependence index, calculated between the current trial and several levels of previous (n-back) trials (x-axis). A positive index indicates a positive serial dependence relationship. The light grey bars represent the serial dependence based on the previous stimulus and the dark grey bars represent the serial dependence based on the previous choice. The data are group means and error bars are 95% confidence intervals and the symbol *** means p < 0.001. **B.** The correlation between participants’ standard deviations of bias across blocks and their artefactual serial dependence on the previous stimulus (open circles, light grey line) and previous choice (filled circles, black line).

Because the grating was present at threshold, the stimulus and choice were highly correlated (75% accuracy). Thus, it is difficult to investigate whether it was the previous stimulus or the previous choice that affected current perceptual decision making. To separate the contribution of stimuli and choices, we examined the influence of one on current perception by controlling the other’s influence. For example, to examine the influence of the previous stimulus, we calculated *c*_*shift*_ for trials preceded by an anticlockwise choice and trials preceded by clockwise choice, respectively, and then averaged these two values. We did the same to calculate the influence of previous choice by averaging the *c*_*shift*_ for trials preceded by clockwise and anticlockwise stimuli. As shown in Figure 3A, we found a positive serial dependence on previous choice for choices made 2-4 trials back from the current trial; 2 back: *M* = 0.11, 95% confidence interval = [0.061 0.16]; 3 back: *M* = 0.059, *CI* = [0.020 0.099]; 4 back: *M* = 0.045, *CI* = [0.0068 0.084]) with one-sample t-tests against zero (2 back: *t*(28) = 4.55, *p* = 9.56e-05; 3 back: *t*(28) = 3.08, *p* = 0.0046; 4 back: *t*(28) = 2.41, *p* = 0.023). The analysis of dependence on the previous stimulus 1-5 trials back revealed no significant influence on current perceptual choices (*t*(28) = −0.27, 0.56, 0.75, −0.28 and 0.79, *ps* = 0.78, 0.58, 0.46, 0.78 and 0.44, respectively. The conclusion that perceptual choices rather than previous stimuli influence current perceptual decision making was also confirmed by comparing the effect of correct choices and incorrect choices. As shown in Figure 3B, a two-way, repeated-measures ANOVA revealed that there was no difference between correct and incorrect choices for 1-5 back analysis (*F*(1, 28) = 0.28, *p* = 0.60). Thus, we demonstrated that the previous percept rather than the previous stimulus *per se* influenced current perceptual decision making.

**Figure 3.**
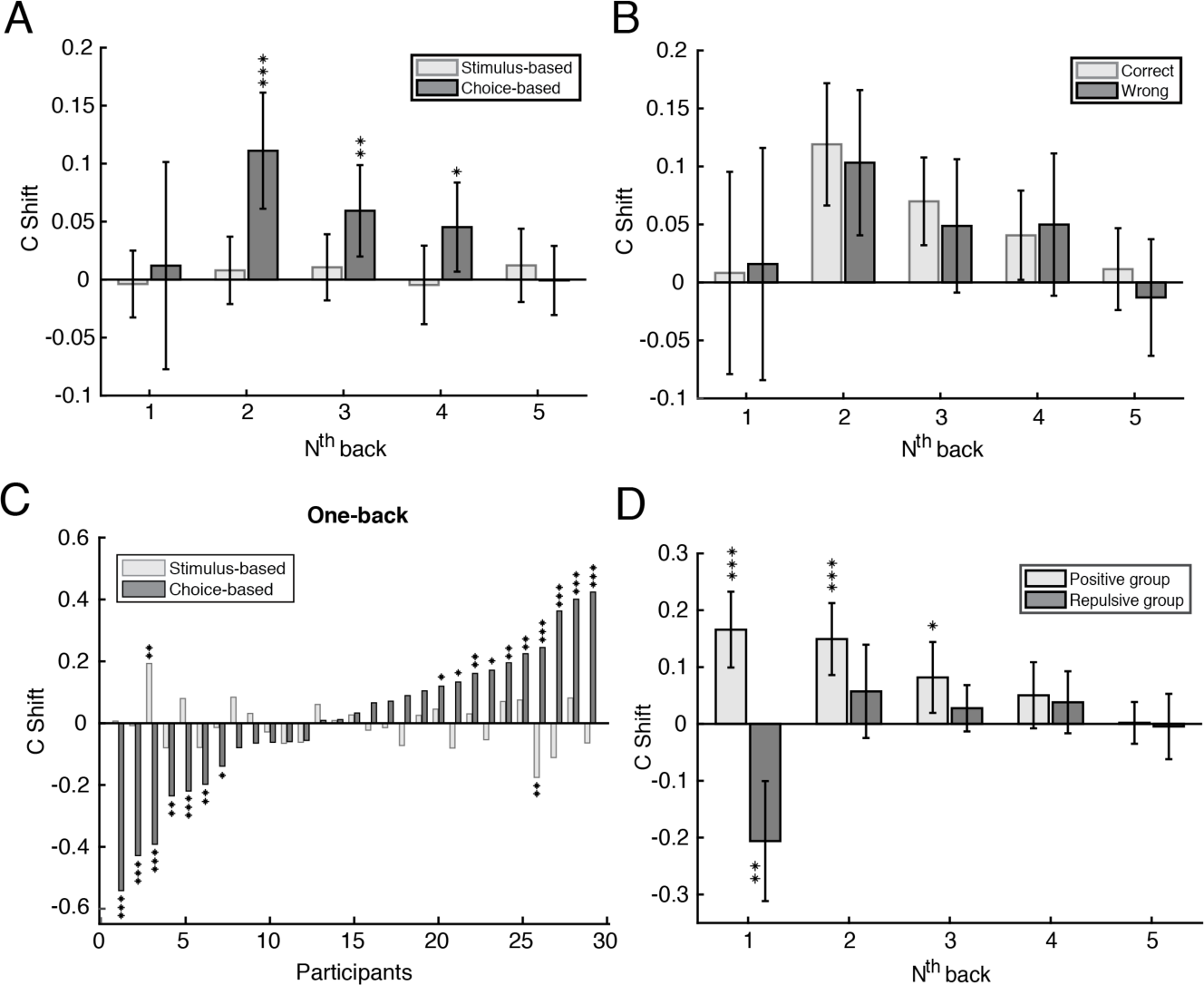
Results from the analysis of serial dependence for Experiment 1. Error bars represent 95% confidence intervals. Symbols ***, ** and * indicate *p* < 0.001, *p* < 0.01 and *p* < 0.05, respectively. **A.** Group mean serial dependence based on the previous stimulus (light grey bars) and choice (dark grey bars). The x-axis indicates the analysis on N trials back from the current trial. The y-axis is the serial dependence index (c shift), with a positive value meaning a positive serial dependence. **B.** The choice data from panel A split by whether the previous choice was correct (light grey bars) or incorrect (dark grey bars). **C.** The one-back serial dependence effect calculated on previous stimulus (light grey bars) and choice (dark grey bars) for each participant. **D.** The choice-based, one-back data from panel C, grouped by whether the sign of serial dependence was positive (n = 17: light grey bars) or negative (n = 12: dark grey bars), showing the persistence of the effects over 1-5 trials back.

Note, the influence of one-back choice (*M* = 0.012, *CI* = [−0.077 0.10]) was not significant (*t*(28) = 0.28, *p* = 0.79), inconsistent with previous serial dependence findings that usually show a largest effect for one trial back (St John-Saaltink el al., 2016; Fischer & Whitney, 2014). Previous perceptual decision-making studies have shown that observers’ biases towards their previous one-back choices can differ markedly among individuals, showing positive or repulsive biases, or even no bias at all (Abrahamyan et al., 2016; Braun et al, 2018). We therefore decided to conduct further analyses to determine whether individual differences caused the non-significant group mean result for the one-back choice-based analysis shown in Figure 1A.

For our analysis of individual participants, we tested for serial dependence based on both previous stimulus and previous choice in each participant’s data using permutation tests. We shuffled the trial sequence within each block and calculated the serial dependence index, *c*_*shift*_. We repeated this procedure 1000 times, producing a null distribution of *c*_*shift*_. The p value was calculated by the proportion of the 1000 *c*_*shift*_ values that were greater than the *c*_*shift*_ value computed with the original data (i.e., one-tailed test) when the original *c*_*shift*_ value was positive. When the original *c*_*shift*_ value was negative, the p value was the proportion of the 1000 *c*_*shift*_ values that were smaller than the original *c*_*shift*_ value. For the choice-based analysis, we found that there were large individual differences, with 10 participants showing significant positive serial effects, 12 participants showing no bias, and 7 showing significant repulsive effects (Figure 3C). In contrast, for serial dependence calculated on previous stimulus, we found significant results for only 2 out of 29 participants, suggesting a consistent non-effect across participants. Furthermore, we looked at the persistence of serial dependence over several levels of n-back by dividing participants into two groups based the sign of *c*_*shift*_ (Figure 3D). For the group showing positive one-back effects, the pattern is similar to traditional serial dependence reports, lasting for 3 trials back with the amplitude decreasing as the number of trials back from the current trial increased (1-3 backs: *M* = 0.17, *CI* = [0.099 0.23]; *M* = 0.15, *CI* = [0.086 0.21]; *M* = 0.082, *CI* = [0.019 0.14]) revealed by one-sample t-tests against zero (1-3 backs: *t*(16) = 5.27, 5.0 and 2.78, respectively; *ps* = 7.6e-05, 1.3e-04 and 0.012, respectively). For the group showing negative serial dependence effects, only the one-back effect (*M* = −0.21, *CI* = [−0.31 −0.10]) was significant, *t*(11) = −4.3, *p* = 0.0013.

These individual differences in choice-based serial dependence are large and even involve effects with opposite signs. What could cause this individual difference? One possibility stems from the fact that the stimulus-response mapping was consistent in Experiment 1, meaning there was an inseparable association between the previous choice and the previous motor response. If the motor response and perceptual decision both generate serial dependencies, but with opposite signs, then the summed effect would be variously positive or negative among observers depending on the relative strengths of each serial component. Experiment 2 tests this possibility by randomising the mapping between stimuli and response buttons.

### Experiment 2

In Experiment 2, we compared the serial effects arising from choices and responses directly. As shown in Figure 1, a cue was presented on the screen after the target was presented to indicate the stimulus-response mapping. There were two types of cue, left click for anticlockwise orientation, right click for clockwise orientation, or the opposite). In the single mapping task, the cue remained the same throughout the experiment, thus similar to Experiment 1. In the random mapping task, the left-right order of the cue lines was randomised on each trial to separate the choice and from the response. All participants finished the random mapping first and then the single mapping task. Since we already showed the previous stimulus *per se* did not influence the following perceptual decision making, here we examined the serial dependence index *c*_*shift*_ for choices and responses. In the single mapping task, the *c*_*shift*_ should be the same for choices and responses because of the consistent stimulus-response mapping for each participant. Thus, we only report the statistics for *c*_*shift*_ computed based on the previous choice (both shown in Figure 4). We removed one participant’s data from the analysis because there was a very large (five-fold) change in the grating’s threshold contrast between the first and second half of the single mapping task (first half, *M* = 0.069, *SD* = 0.011; second half, *M* = 0.33, *SD* = 0.13), suggesting very poor performance in the second half.

**Figure 4.**
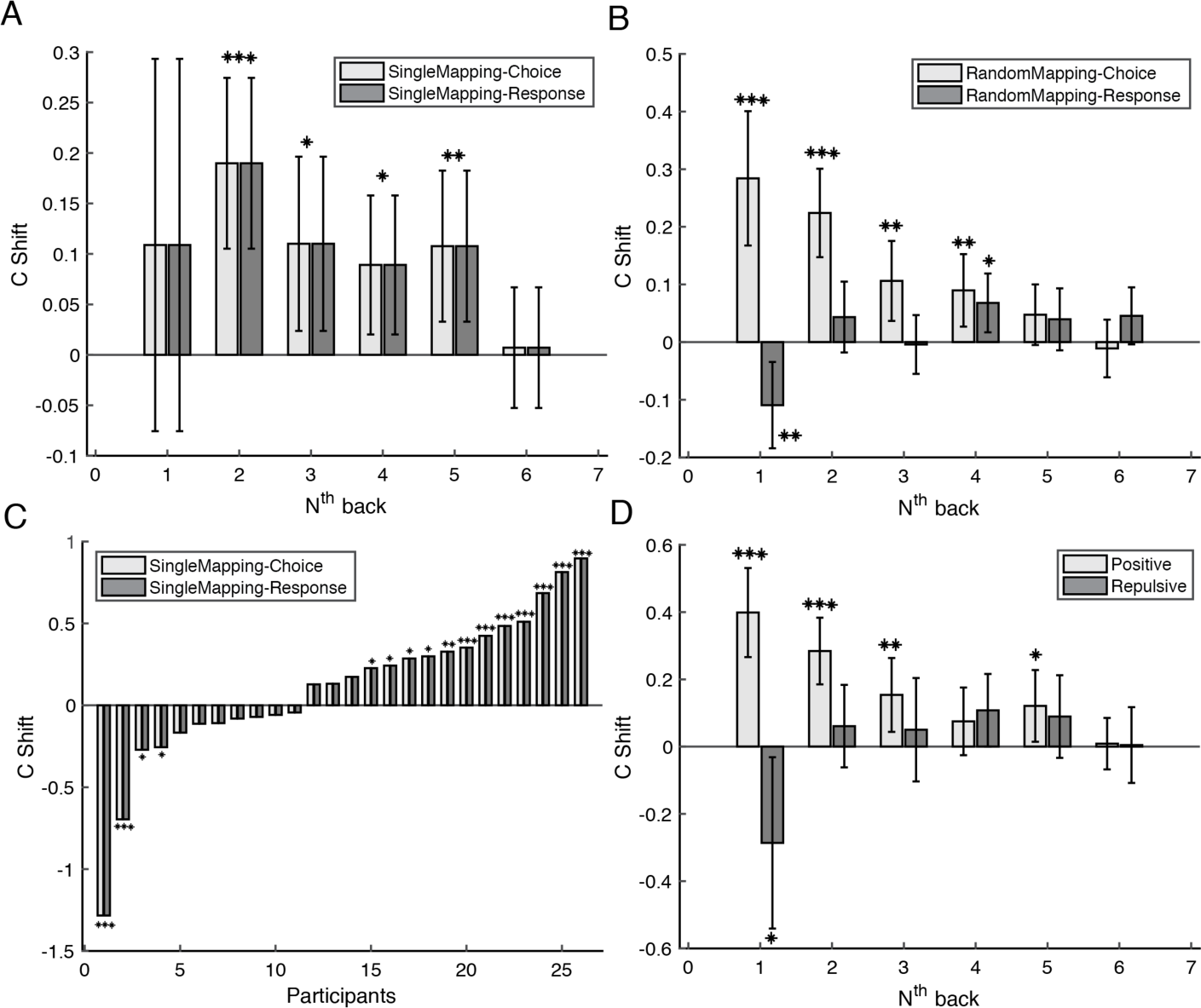
The analysis results on serial dependence for experiment 2. Error bars represent 95% confidence intervals. Symbols, ***, **, and *, mean *p* < 0.001, *p* < 0.01, and *p* < 0.05, respectively. **A.** The serial dependence on previous choice (light gray bars) and response (dark gray bars) in the single mapping task. The x axis indicates the analysis on N trials back from the current trial. The y axis is the serial dependence index (C shift), with a positive value meaning positive serial dependence. **B.** The serial dependence on previous choice (light gray bars) and response (dark gray bars) in the random mapping task. The x axis indicates the analysis on N trials back from the current trial. The y axis is the serial dependence index (C shift). **C.** Serial dependence on previous one-back choice (light gray bars) and response (dark gray bars) for each participant in the single mapping task. **D.** Persistence of serial dependence for two groups of participants in the single mapping task: one with positive serial dependence on the one-back choice (light gray bars) and the other with negative serial dependence on the one-abck choice (dark gray bars).

For the single mapping task (similar to Experiment 1), we found a similar pattern of serial dependence on previous choice/response (Figure 4A, compare with Figure 3A). The one-back effect was not significant (*t*(25) = 1.2, *p* = 0.24), but a positive serial dependence was found for 2-5 trials back: *t*(25) = 4.6, 2.6, 2.7 and 3.0, respectively; *p* = 1.0e-04, 0.015, 0.013 and 0.0067, respectively. For the random mapping task, where the perceptual choice and motor responses were separable, a typical serial dependence on previous choice was found (Figure 4B). The serial dependence persisted 4 trials back with the amplitude decreasing as the n-back interval increased (1-4 trials back: *M* = 0.28, 0.22. 0.10 and 0.090, respectively; *CI* = [0.16 0.40], [0.15 0.30], [0.04 0.18] and [0.027 0.15], respectively), revealed by one-sample t-tests (*t*(25) = 5.0, 6.0, 3.2 and 2.9, respectively; *p* = 3.5e-05, 2.8e-06, 0.004 and 0.007, respectively). As shown in Figure 4B, we found a repulsive serial dependence on previous motor response (*M* = −0.11; *CI* = [−0.18 −0.035]) for one trial back (*t*(25) = −3.0; *p* = 0.0059). There was a positive effect (*M* = 0.068; *CI* = [0.017 0.12]) for analysis on four trials back (*t*(25) = 2.75; *p* = 0.011).

We also looked at the individual differences for one-back serial dependence for the single mapping task, just as we did in Experiment 1. By performing a permutation test on each participant’s data, we showed that participants variously had a positive bias (N = 12), a repulsive bias (N = 4) or no bias at all (N = 10) towards the previous choice/response (Figure 4C). Note, the same group of people showed a consistent positive bias towards one-trial back choice in the random mapping task. We divided participants into two groups based on the sign of their one-back serial dependence (negative or positive *c*_*shift*_) to check the persistence of serial dependence over greater n-back distances (Figure 4D). For people with a positive one-back effect, the amplitude of serial dependence decreased as the temporal distance increased (1-3 back: *M* = 0.39, 0.28 and 0.15, respectively; *CI* = [0.27 0.53], [0.19 0.38] and [0.044 0.26], respectively) revealed by one-sample t-tests (*t*(14) = 6.5, 6.2 and 3.0; *p* = 1.5e-05, 2.5e-05 and 0.0095). For people showing a repulsive one-back effect (*M* = −0.28, *CI* = [−0.54 −0.032]; *t*(10) = −2.5, *p* = 0.031), they showed a positive effect for analysis on four trials back (*t*(10) = 2.22, *p* = 0.05). Thus, the patterns of two groups of participants with either positive or negative one-back serial dependences (Figure 4D) was similar to the serial dependence on the previous choice or previous motor response in the random mapping task. This suggests that the individual differences for one-back choice/response we observed in Experiment 1, and in the single mapping task of Experiment 2, reflect the different weighted average of positive bias for perceptual choices and repulsive bias for motor responses.

## Discussion

In two experiments, the current study examined serial dependence in orientation discrimination under conditions of stimulus uncertainty (contrast controlled at threshold level). By well-controlled data analysis and manipulation of stimulus-response mapping, we clarified the roles of stimulus, perceptual choice, and motor response in serial effects. Our results showed that the physical stimulus *per se* did not influence subsequent perceptual decision making but that the percept of the stimulus did affect subsequent perceptual choices in an attractive way. In addition, we found that the motor response exhibited a negative serial dependence, being repelled away from the preceding motor response. Moreover, when the choice-response contingencies were consistent and thus inseparable, individual differences in overall serial effect was observed, which was likely due to different observers giving different relative weights to the positive bias for perceptual choice and the repulsive bias for motor response. The clarification of different serial dependences for perceptual choice and motor response may help resolve some contradictions in reported findings, particularly in categorization tasks, about whether serial dependence is positive or negative.

Consistent with previous findings (Fründ et al., 2014; St John-Saaltink et al., 2016), the current study confirmed that the percept of the stimulus instead of the physical stimulus *per se* influenced subsequent perception in an attractive way. With continuous perceptual report measures, such as orientation reproduction, it has been suggested that serial dependence operates on perception (Cicchini et al. 2017; Fischer & Whitney, 2014) and decision (Fritsche et al., 2017). The current study differs from these studies in two aspects. First, in contrast to continuous measures, the task here is a binary forced choice between two discrete stimuli (45° and −45° orientations). Second, the stimuli were embedded in noise with grating contrast controlled at threshold level and the duration of the stimulus was short (6.3 ms compared with hundreds of ms in reproduction tasks). Thus, unlike the stimulus presented in the reproduction task, which would induce a salient and clear perception of orientation in a fine-grained range, perception of stimuli in the current study has a much larger uncertainty. Serial dependence may differ between these different tasks. When examining the serial effect using a reproduction task, the previous stimulus can be used as the indicator of the percept because the percept of stimulus is well represented by the physical stimulus (a 45° orientation is rarely perceived as − 45°) and motor errors are involved in the reproduction. For a binary categorization task under uncertain conditions, such as used here, the percept might dramatically differ from the physical stimulus *per se* on a given trial. Thus, the response should be used to represent percept when examining the serial dependence.

What is the neural mechanism underling current serial dependence on previous choices? Using fMRI, St John-Salltink et al. (2016) showed that the bias towards previous percept is reflected in activities in primary visual cortex. The serial dependence on previous choice found in the current study is likely a perceptual effect (Cicchini et al. 2017; Fischer & Whitney, 2014). However, it may reflect the influence of top-down expectations rather than bottom-up accumulation of sensory evidence (no influence from stimulus *per se* was observed). In fact, studies have shown that prior expectations can bias sensory representations in the visual cortex (Kok, Failing, & de Lange, 2014; Summerfield & de Lange, 2014). Moreover, these prior expectations induce the preactivation of stimulus templates before stimulus onset (Kok, Mostert, & de Lange, 2017). Since natural visual statistics are dominated by slow-changing components (Dong & Atick 1995), thus predicting temporal correlation over short period, exploiting this prior acquired over long-term life experience is a very sensible strategy for achieving a stable and accurate perception of the world, especially given that the sensory evidence is inevitably noisy. The positive serial dependence we found in the current study may reflect the usage of long-term prior expectations on a relatively stable world even though trials presenting different stimuli were randomized in our laboratory task.

The finding of a repulsive serial effect on motor responses supports the idea that action is more than a final output stage after a decision has been made and instead is actively involved in response selection during sensorimotor decision making (Cisek & Kalaska, 2005; Klaes et al., 2011; Pape & Siegel, 2016; Pastor-Bernier & Cisek, 2011). For the task with a random stimulus-response mapping in Experiment 2, the effects of choice and motor response were separated. Although the task was to judge which orientation was presented for a given trial, the ultimate goal was to execute an action. The alternation bias on motor response we found here is consistent with previous studies (Pape et al., 2017; Pape & Siegel, 2016). Notably, in Pape and colleagues’ studies (2016, 2017), observers used left and right index fingers to respond, while in the current study, index and middle fingers of the right hand were used. Thus, this alternation bias on motor response may generalize to different forms of binary movement involving response competition (e.g., binary saccadic eye movement). Using an experimental design with a consistent choice-response contingency, previous studies found that neural activities over motor areas predicted choice bias (and response bias, too, as stimulus and response were correlated) (de Lange, Rahnev, Donner, & Lau, 2013; Donner, Siegel, Fries, & Engel, 2009). By separating the choice content and motor response, Pape and Siegel (2016) showed that beta-band (12-30 Hz) activities in motor cortex predicted response alternation. Together with previous studies, the repulsive serial effect on motor response suggests that during perceptual decision making, the final action is not only a faithful output of perceptual choice, even if the task setting encouraged a sequential process. In other words, the motor system itself also actively contributes to response selection.

Why is the serial dependence on motor response repulsive? One possibility is because of motor efforts. It has been proposed that motor control is decision making (Wolpert & Landy, 2012), which can be influenced by motor effort (Cos, Belanger, & Cisek, 2011; kibbe & Kowler, 2011). Moreover, perceptual decisions were also found to be influenced by the cost to act — perceptual choices linked to energetically more-costly motor responses were avoided (de Lange & Fritsche, 2017; Hagura, Haggard, & Diedrichsen, 2017). In the current study, it is possible that repetitively clicking the same button caused muscle fatigue, especially during a long testing session (a one-hour session for the task with random stimulus-response mapping). The alternating motor bias may help in this case to reduce motor fatigue. Another possibility is that the repulsive bias results from the exploratory nature of the action. For example, when actively searching for an object in the environment, we voluntarily saccade away from the region that was previously fixed, a phenomenon known as inhibition of return in visual search (Klein & Maclnnes, 1999). To initiate a different action to explore the outside world is beneficial after a failure of the previous motor exploration (e.g., more likely to find the target). Because the perceptual decision making in the current study was under conditions of uncertainty, and observers did not receive any feedback regarding ‘right’ or ‘wrong’ responses, the observers were likely to switch responses after an unsure response.

As used in many studies, the stimulus-response mapping is often consistent during the task, which causes individual differences on serial dependence. This is because the overall serial dependence is the weighted average of positive serial dependence from perceptual choice and repulsive serial dependence from motor response. Different observers have different inherent preferences for these two types of serial dependence. Future studies can further examine whether these individual preferences are related to other intrinsic biases, e.g., the exploration of different options versus the exploitation of their reward (Hills, Todd, Lazer, & Redish, 2015; Mehlhorn et al, 2015). Previous studies have shown that these inherent positive or repulsive serial dependence biases can adapt to the temporal statistics of the task (Abrahamyan et al., 2016; Braun et al., 2018), suggesting a flexible mechanism underling the overall serial bias. In previous studies using consistent stimulus-response mappings, some found repulsive serial dependence (Fründ et al. 2014; Taubert et al, 2016). It is not impossible that this reflects the repulsive motor bias rather than a true repulsive perceptual effect. The current findings of individual differences in serial dependence and its two opposite components (positive serial dependence on perceptual choice and repulsive serial dependence on motor response) highlight that careful attention must be given to careful experimental designs when examining serial dependence.

To conclude, the current study shows that serial dependence in perceptual decision making under conditions of uncertainty operates on perceptual choice and motor response in opposite directions, a positive bias for previous perceptual choice and a repulsive bias for previous motor response. We did not find any influence of the physical stimulus itself on subsequent perception, suggesting the positive bias towards previous perceptual choice may reflect the top-down influence of expectation of temporal continuity of the visual world. When choice-response contingency is held constant, the relative weights of the positive perceptual bias and the repulsive motor bias varies among individuals, meaning that the overall effect of serial dependence (i.e., positive, repulsive or no bias) shows considerable individual differences. The current study thus elucidates a key reason for some of the contradictions reported in the field of serial dependence regarding the sign of the serial effect.

## Acknowledgements

This work was supported by Australian Research Council grant DP150101731 to DA. We thank Professor David Burr and Dr. Guido Marco Cicchini for constructive suggestions concerning data analysis.

